# Comparative Anatomical, Ultrasonographical and Radiological studies of the biliary system of Rabbits and Domestic cats in Egypt

**DOI:** 10.1101/834408

**Authors:** R.T. Reem, M.A. Maher, H.E. Alaa, H.A. Farghali

## Abstract

Under the prevailing overall Conditions of all veterinarians for the diagnosis of biliary diseases, application of surgical procedures and liver transplantation in Cats as carnivorous pet animal, and Rabbits as herbivorous pet animal and also as a human model in research. The present study was constructed on twelve native breeds of rabbits (Oryctolagus cuniculus) and eighteen adult domestic cats (Felis catus domesticus). We concluded that, in brief; the rabbit gall bladder was relatively small, fixed by several small hepato-cystic ducts to its fossa. The rabbit bile duct was formed commonly by the junction of the left hepatic duct and the cystic duct. The cystic duct was commonly fairly large, received the right hepatic duct that collected the right lobe in its route to enter the duodenum, the bile duct receives the branch of the caudate process of the caudate lobe. The present study revealed other four anatomic variations dealing with the shape and size of the feline native breed’s gall bladder from fundic duplication, bilobed, truncated fundus and distended rounded fundus. Commonly, the bile duct was formed by the triple convergence of the left and the right hepatic ducts with the cystic duct. However, in some exceptional cases a short common hepatic duct was formed. Sonographically, the normal gall bladder in rabbit appeared small, elongated with anechoic lumen bordered by right lobe laterally and quadrate lobe medially and has no visible wall, but in cat varied in conformation, bordered by the right medial lobe laterally and the quadrate lobe medially surrounded by echogenic wall.

## INTRODUCTION

The biliary system comprised; the gall bladder, cystic duct, bile duct and hepatic ducts with its intrahepatic ramifications **[1, 2].**

The common hepatic duct was missing in rabbits, cats and dogs. In rabbits, two hepatic ducts were described; the left one drained both the left lobes and the quadrate lobe, then joined the cystic duct to form the bile duct which received the right duct, the latter drained the right and the caudate lobes. However, in feline and canine, the lobar hepatic ducts opened individually into the cystic duct **[3].** In this respect, the absence of the common hepatic duct was also recorded in rabbit **[4]** and guinea pig **[5] [6].** The common hepatic duct was formed in the rabbit by the confluence of the left and the right hepatic ducts at the ventral part of the liver porta. It received ramus omentalis and at the same level the cystic duct to form the bile duct. The latter duct received the ramus processus caudati through its course in the hepatoduodenal ligament **[7].**

In cats and dogs, 3-5 hepatic ducts opened separately in to the cystic duct which became the bile duct after the entrance of the last hepatic duct **[8]; [9]; [10] and [11].** On the other hand, the cystic duct extended from the gall bladder to the first hepatic duct **[1].** However, the cystic duct is an important landmark in that it distinguishes the otherwise continuous hepatic ducts from the bile ducts **[2].** In cats, the bile duct was formed by the triple convergence of the left hepatic, the right hepatic and the cystic ducts **[12]**.

In ruminants, the bile duct was formed by the union of the right and the left hepatic ducts, while the cystic duct opened into the right duct **[13]; [14] and [15].** On the other hand, the right and left hepatic ducts joined to form a short common hepatic duct, which received the cystic duct and then after became the short and wide bile duct **[16]; [10] and [17].**

The rabbit gall bladder was long oval **[7]**, cylindrical sac **[3]** or had elongated body with narrow neck **[4]**. It was embedded in its deep fossa at the middle of the visceral surface between right and quadrate lobes. It did not reach the liver ventral margin even when it was completely filled.

In carnivores, the gall bladder was sunk deeply between the right medial and quadrate lobes, just to the right of the median plane and opposite the eighth intercostal space. It was visible on the visceral as well as the parietal (diaphragmatic) surfaces and thus, in contact with the diaphragm **[10] and [18].** However, the guinea pig gall bladder was rounded and extended to the liver ventral border, hanging freely and suspended by a single membrane from the liver **[19].** Also, and in this respect, the pear-shaped gall bladder of ruminants was firmly attached to its fossa on the liver visceral surface, and protruded from its ventral margin **[10].**

The majority of the congenital abnormalities of feline gall bladder could be related to the failure of vacuolization of the solid gall bladder and bile duct diverticulum **[20].** Three types of gall bladder abnormalities were recorded; a. Septated gall bladder, had a large lumen divided by septa inside, b. Bilobed gall bladder, with fundic duplication, body duplication where the two lobes opened by a single cystic duct to the bile duct, c. Duplex gall bladder, where the fundus, the body and cystic duct were formed as a couple (complete duplication) and they joined together at the bile duct **[21]; [22] and [23].** Also in feline, a case report in anorexic cat with distended rounded gall bladder at the ventral hepatic margin was cited **[24]**.

The gall bladder was lacking in the horse **[25]; [16] and [10]**, in camel **[26]**, in deer **[27]** and in rat **[28]** which was compensated by enlargement of the duct system **[18].**

Ultrasonography is an excellent imaging tool in the diagnosis of gall bladder abnormalities **[29]**. It’s the imaging technique of choice in veterinary medicine to localize the gall bladder easily in the right cranial abdomen **[2]**. The gall bladder and cystic duct can be easily identified ultrasonographically in clinically normal cats, however the remaining extrahepatic and intrahepatic structures are not visible unless there is pathological condition **[30] and [31]**.

## MATERIALS & METHOD

The present study was constructed on twelve (12) native breeds of rabbits (*Oryctolagus cuniculus*) and eighteen (18) domestic cats (*Felis catus domesticus*) which were adult apparently healthy of both sexes with average body weight; 2.800-3.500 Kg.

### A. The Anatomical study

The used animals were euthanized by lethal dose of Diazepam at 10 mg/kg intravenously through the external jugular vein. The experiments were conducted by the international ethical standards set by **the institutional animal care and use committee (Vet. CU. IACUC) VetCU1111201808**. Three rabbits and nine cats were anatomically dissected freshly for gall bladder morphology then fixed by 10% neutral buffered formalin. For cast study, six rabbits and nine cats were injected with colored latex neoprene into the major duodenal papilla and then fixed by freezing or neutral buffered formalin overnight, further dissection occurred and then transferred to concentrated HCL and left for three days.

The specimens were photographed using Olympus digital camera SP-600UZ 12 mega pixel. The anatomical nomenclature used in this study was in accordance with the ***Nomina Anatomica Veterinaria 2017*** (6^th^ edition).

### B. The Ultrasonographical and Radiological study

Concerning the Ultrasound examination, we evaluated 6 live animals; 3 each animal species, by using two devices according to availability at the time of study; B-mode scan (Pie medical) and doppler device (EXAGO, Echo control medical, France) with a linear multifrequency transducer using a frequency of 5-7.5 MHz in rabbits and 7.5 MHz in cats. These animals were later used as specimens for the radiological examination of the biliary duct system using the radiographic device (Fisher imaging, Chicago, USA). The animals were euthanized as mentioned before and injected by lead oxide dissolved in turpentine oil into the major duodenal papilla in both animals to obtain X-ray films.

## RESULTS

### A. Anatomical study

The present investigation in the rabbit, showed a relatively small and elongated gall bladder (Figs 1,2,3/f) with nearly rounded fundus and narrow neck. It was situated and fixed in the gall bladder fossa on the middle of the liver visceral surface, bordered medially by the quadrate lobe and laterally by the right lobe. several small ducts entered the fundus and the corpus of the gall bladder as hepatocystic ducts, were demonstrated during the dissection process of the injected specimens. In addition to their drainage to the parenchyma of the gall bladder fossa, they served as a means of fixation in its highest position and did not permit it to reach the liver ventral margin (Fig. 1, 2A,B/f). It must be noted that the common hepatic duct was missing in the most (common) of rabbit liver specimens. The bile duct (Figs. 2,3f/1) was formed ventral and slightly to the left of the hepatic porta by the junction of the left hepatic duct (Figs. 2,3f/3) and the cystic duct (Figs. 2,3f/5).

**Fig. (1):**
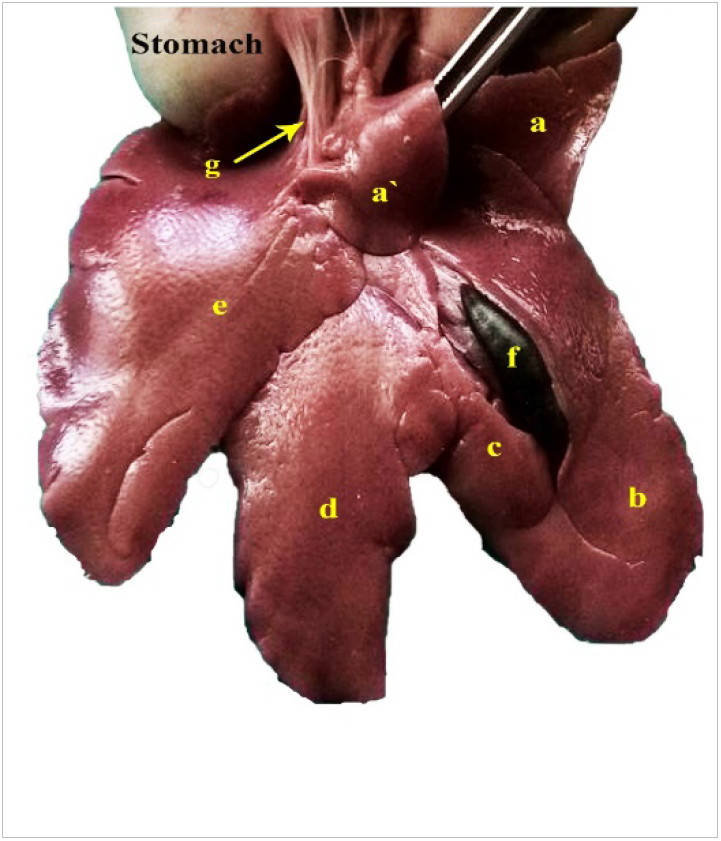
A photograph showing the position of the gall bladder on the visceral surface of the rabbit’s liver

**Fig. (2):**
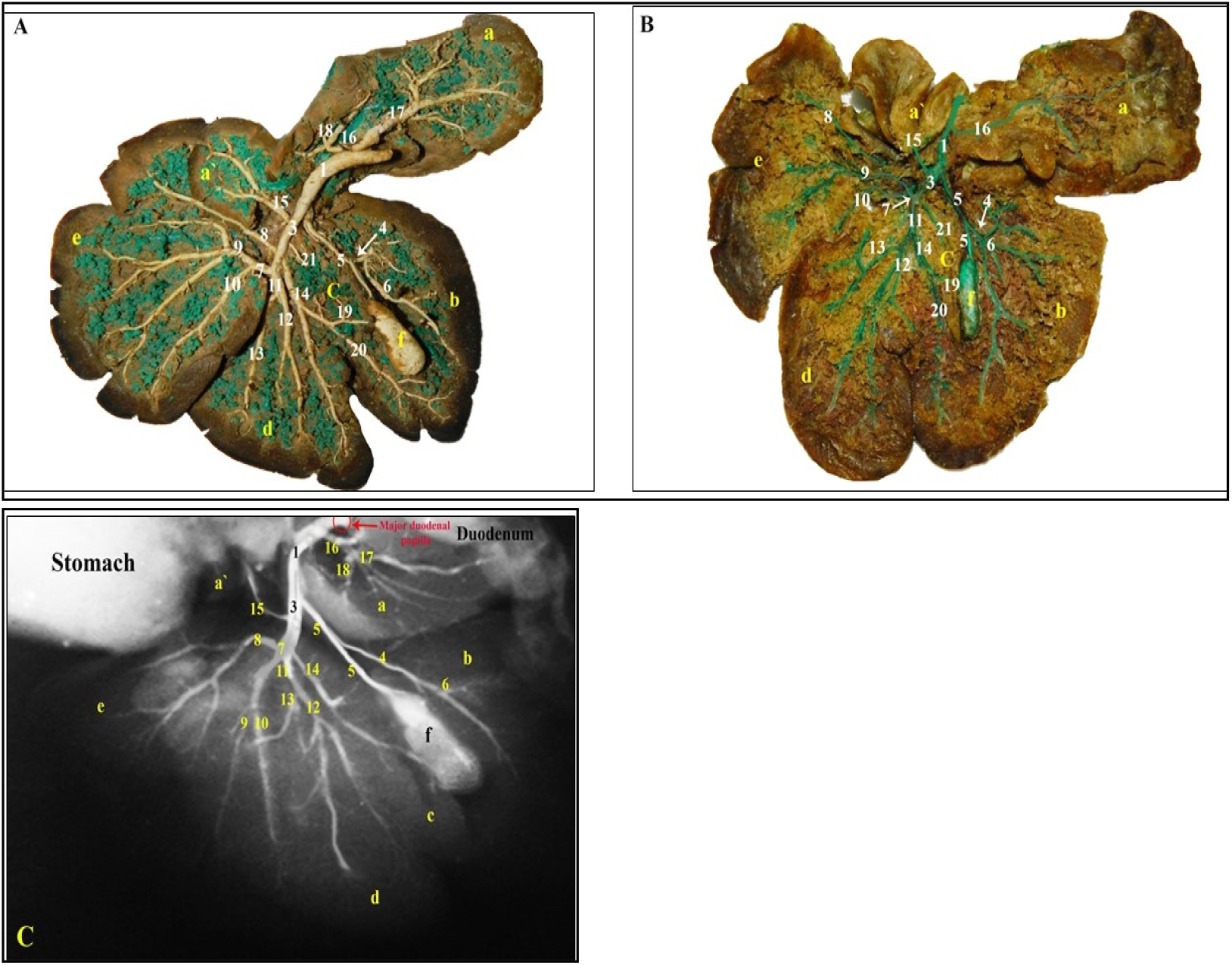
Visceral surface of the rabbit liver showing the biliary system; (A, B): injected with latex neoprene, (C): X-ray film.

**Fig. (3):**
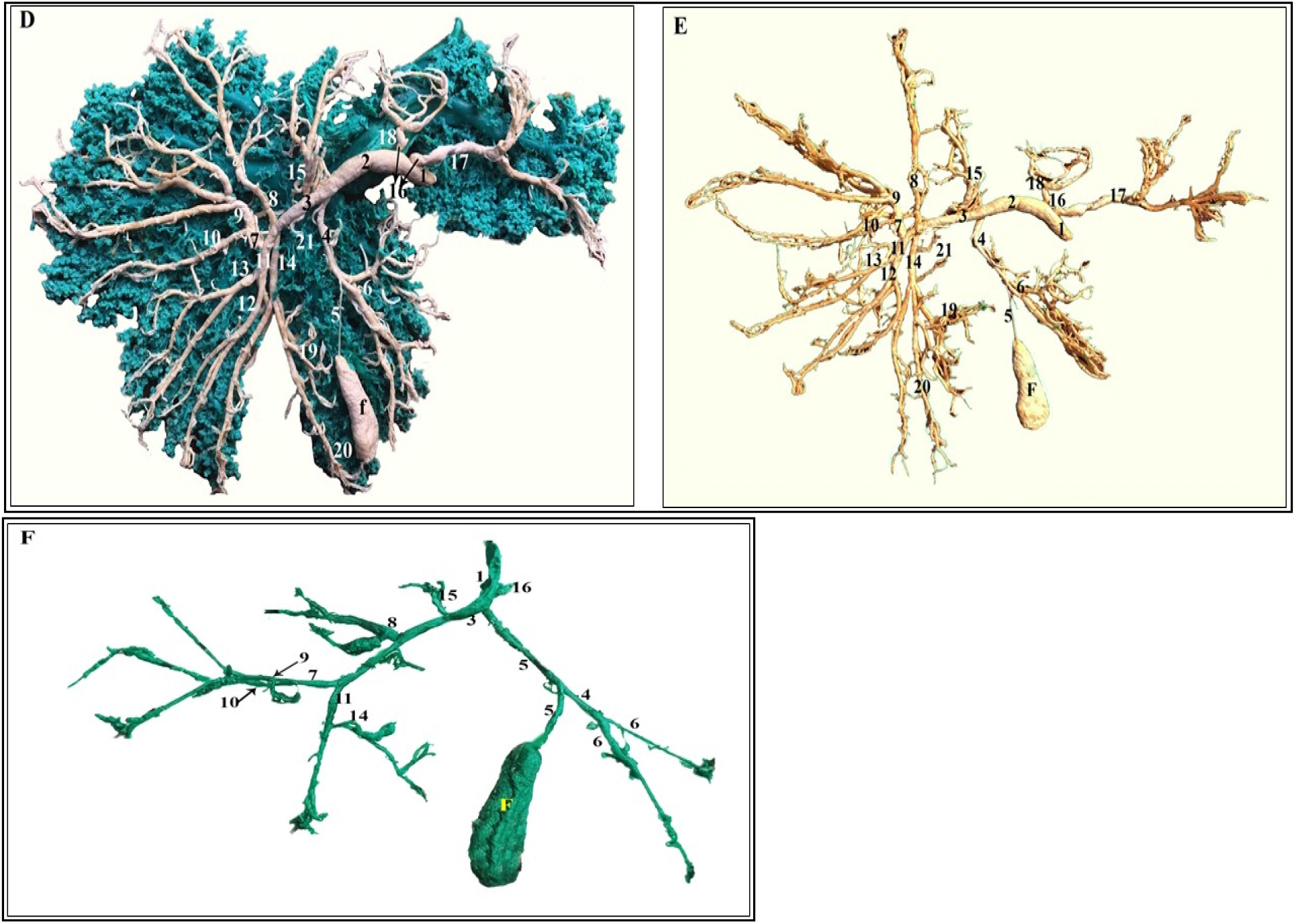
Latex neoprene cast of the rabbit biliary system; (D): biliary system ‘white’ with the hepatic veins ramifications ‘blue’, (E): Latex neoprene cast of the rabbit biliary system isolated ‘white’, (F): Latex neoprene cast of the gall bladder, the terminations of the cystic duct and hepatic ducts and the beginning of the bile duct isolated ‘green’.

#### Ductus hepaticus sinister

(Figs. 2,3/3) drained bile from both left lobes, quadrate lobe and the papillary process of the caudate lobe. It was formed on a level with the umbilical fissure, by the union of the left lateral lobar duct (Figs. 2,3/7) and the left medial lobar one (Figs. 2,3/11). The former duct was short and branches into dorsal, intermediate and ventral radicles (which accompanied the corresponding portal venous branches and its subsequent divisions through the parenchyma of the left lateral lobe (Figs. 2B, C/7,8,9,10), whereas in some cases the dorsal branch arose directly from the parent duct; the left hepatic duct (Figs. 2A, 3/ 7,8). The larger latter duct (Figs. 2,3/11) coursed ventrad following the pars umbilicus venae portae and its ramifications to drain the left medial lobe via 3-4 branches (Figs. 2,3D,E/ 12,13) as well as the quadrate lobe through 2-3 rami (Figs. 2B,3F/ 14, 19, 21). However, the quadrate lobe found to be drained immediately into the parent left hepatic duct in some of the examined specimens (Figs. 2A,C & 3D,E/14,19,21). Beyond its formation, the left hepatic duct (Figs. 2,3/ 3) proceeded along the cranioventral aspect of the pars transversa venae portae where it received the R. omentalis (Figs. 2,3/15) from the papillary process of the caudate lobe, in addition to several twigs from the surrounding hepatic parenchyma. It terminated shortly before the ventral aspect of the liver porta by joining the cystic duct forming the bile duct.

#### Ductus cysticus

(Figs. 2, 3F/5), in the majority of the examined rabbit liver, it was straight, relatively thick and travelled dorsad for about 2.5-3 cm to join the left hepatic duct. Along its course it received several small ducts from the dorsal portions of both right and quadrate lobes, in addition to the right hepatic duct (Figs. 2, 3F/4). **Ductus hepaticus dexter** (Figs. 2, 3F/4) was represented by a short trunk, opened into the right aspect of the cystic duct, about its middle. It was divided into dorsal, intermediate and ventral radicles (Figs. 2,3/6) which accompanied their respective portal venous branches to drain the different portions of the right hepatic lobe. In two out of six examined rabbit livers, it was observed that the cystic duct (Figs. 3D,E/5) was very thin and small connected the gall bladder neck to the fairly large right hepatic duct (Figs. 3D,E/ 4). The latter accompanied its satellite portal venous branches, drained the different portions of the right lobe and proceeded dorsally to join the left hepatic duct (3D,E/3) to form a short common hepatic duct (3D,E/2). The latter, after receiving a fairly large R. processus caudatus from the caudate process and isthmus of the caudate lobe (Figs. 3D,E/ 16), continued as the bile duct (Figs. 3D,E/1).

#### R. processus caudatus

(Figs. 2, 3/16): was fairly large drained its respective large process of the caudate lobe through two radicles (Figs. 2A,C & 3D,E/17,18) that accompanied the satellite portal venous branches and its ramifications. They united to pierce the caudate process root and opened directly into the right aspect of the continuation of the common hepatic duct as the bile duct (Figs. 3D,E/16) or directly into the bile duct (Figs. 2, 3F/16), on a level with the caudate process isthmus. **Ductus choledochus** (Figs. 2,3,4/1): from the point of its formation the bile duct proceeded dorsolaterally through the hepatoduodenal ligament for a short course, about 1.5-2 cm (Fig. 4A/h), to enter the duodenal wall and opened separately on the major duodenal papilla (Figs. 2C, 4B) about 1 cm distal to the pylorus.

**Fig. (4):**
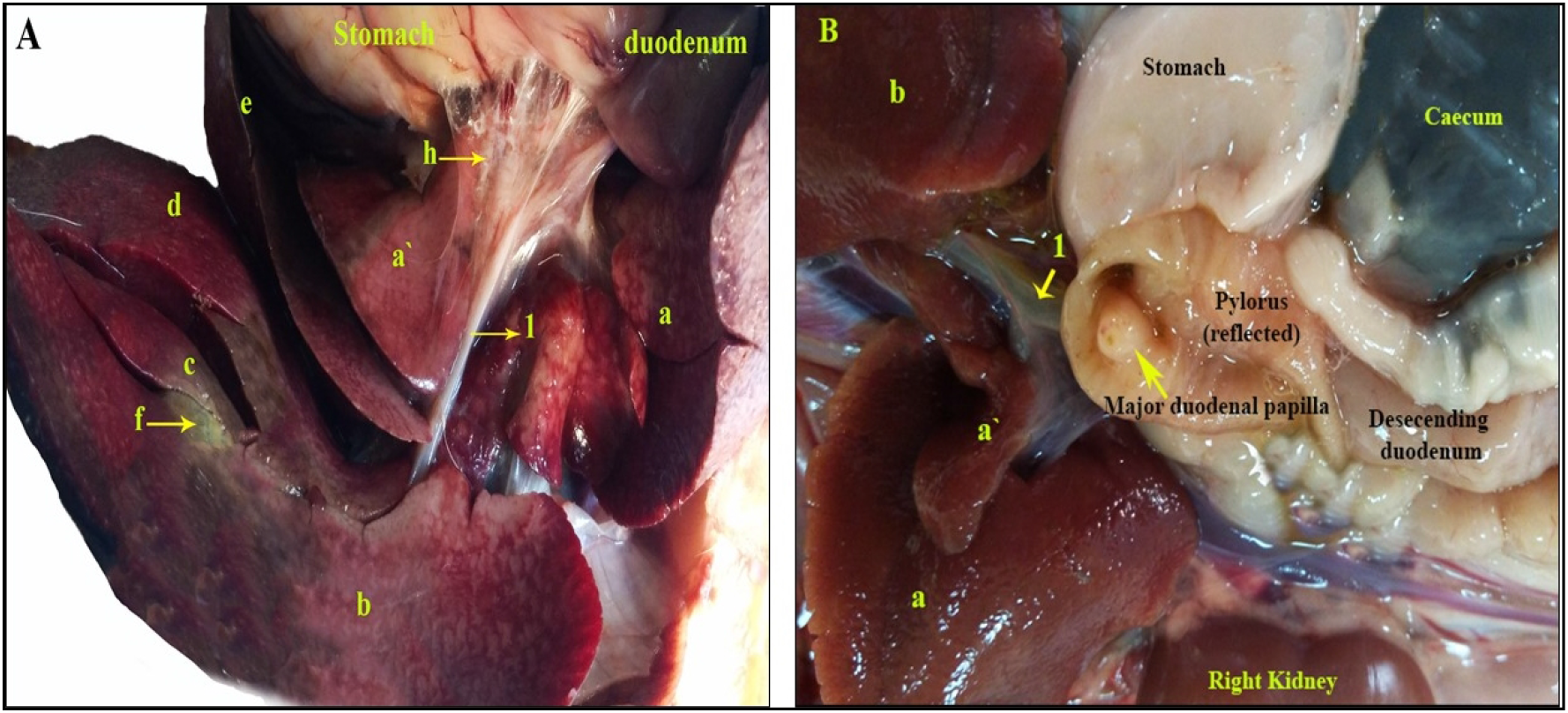
A photograph showing the course of the bile duct in the hepatoduodenal ligament of rabbit (A) and its opening on the major duodenal papilla (B).

It was worthy to note that the biliary system of the native breed cats presented considerable variations in size, shape and position of the gall bladder as well as the manner of terminations of the hepatic ducts and the formations of the bile duct. In the most common investigated cat livers, the gall bladder was relatively large, situated more superficially in the gall bladder fossa, between the right medial lobe laterally and the quadrate lobe medially and did not reach the liver ventral margin and did not appear on the hepatic parietal surface (Figs. 5A,B,C&11A/g). It was curved pear-shaped with broad fundus, slightly concave hepatic surface, slightly convex gastric surface and folded neck (Figs. 5D,E&9B/g) connected with tortuous cystic duct. In two investigated liver specimens, the gall bladder appeared oval in shape (Fig.5F/g). In some of the examined specimens, the gall bladder appeared on the parietal surface of the liver, positioned at the level of the 7^th^ intercostal space (Figs. 6, 7C & 8B,E/g).

**Fig. (5):**
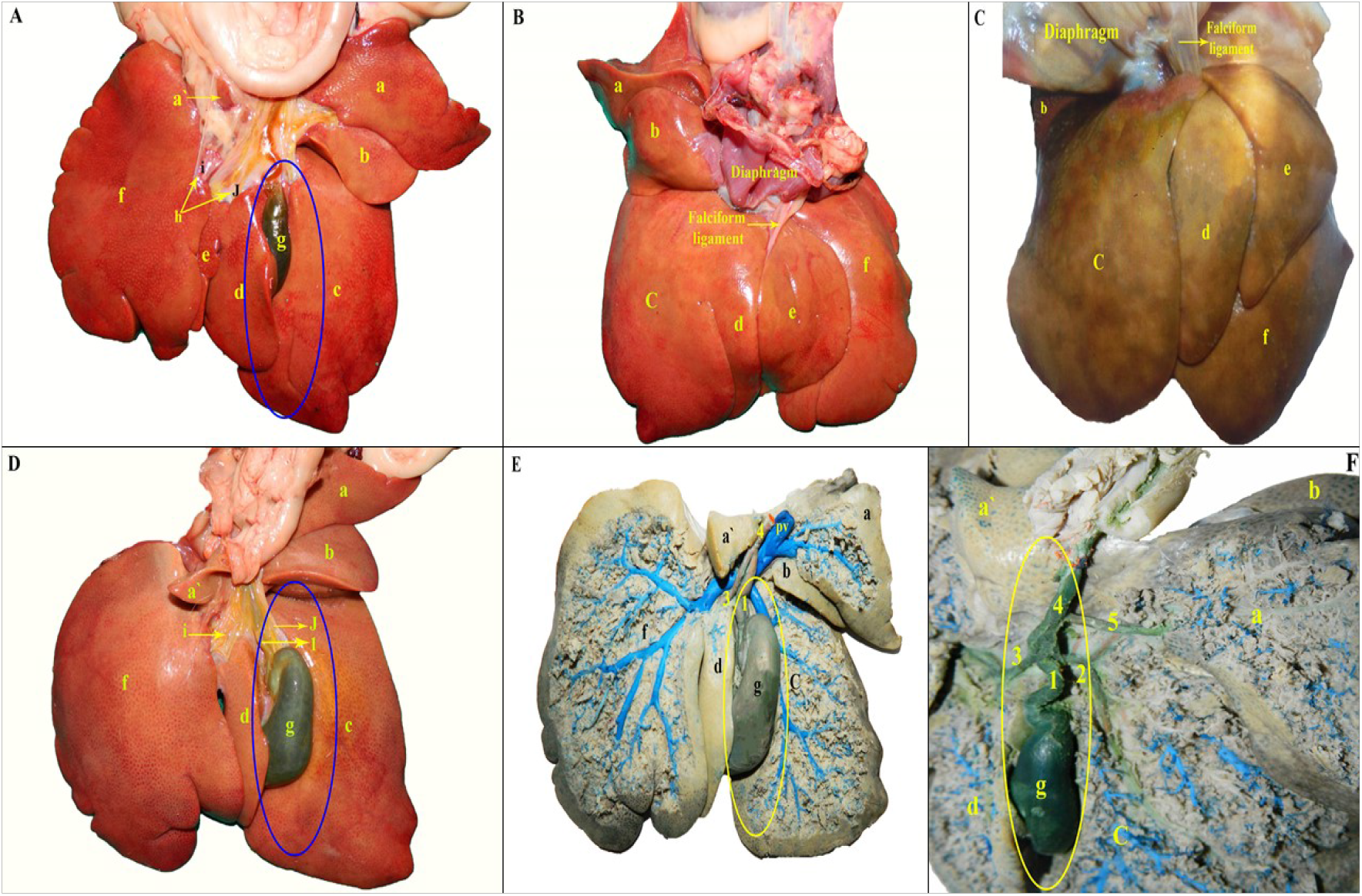
A photograph showing the position of the gall bladder on the Visceral surface of the cat’s liver. (A): Superficially in the gall bladder fossa, between the right medial and the quadrate lobes not reaching the liver ventral margin, (B, C): Not appearing on the hepatic parietal surface, (D,E): Curved pear-shaped gall bladder with tortuous cystic duct, (F): Oval in shape gall bladder.

**Fig. (6):**
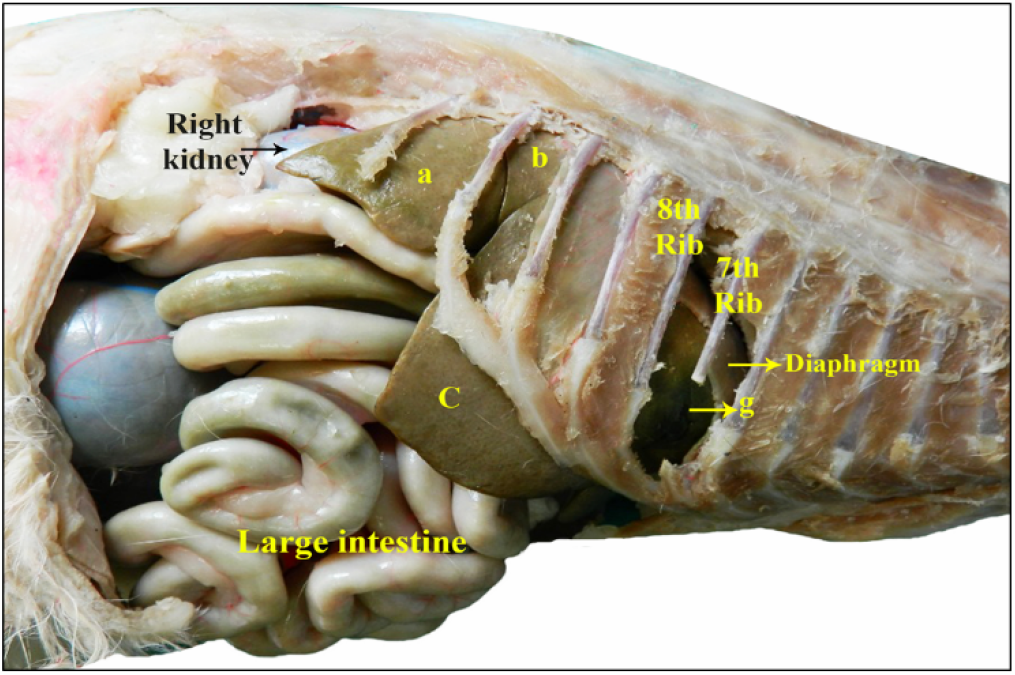
A photograph showing the position of the gall bladder on the parietal surface of the liver of cat at the level of the 7^th^ intercostal space.

**Fig. (7):**
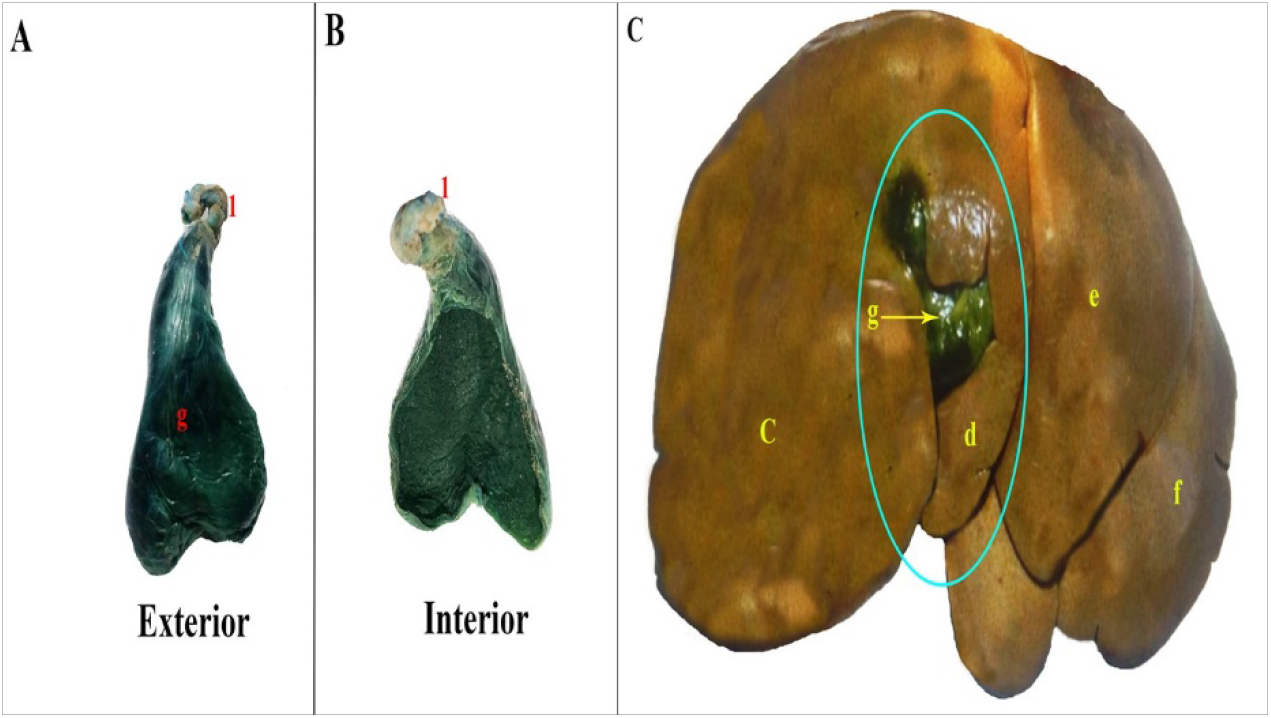
A photograph showing the duplex fundus gall bladder in cat. (A): divided by a sagittal groove on the outside, (B): No internal septa, (C): Duplex fundus gall bladder on the parietal surface of the liver.

**Fig. (8):**
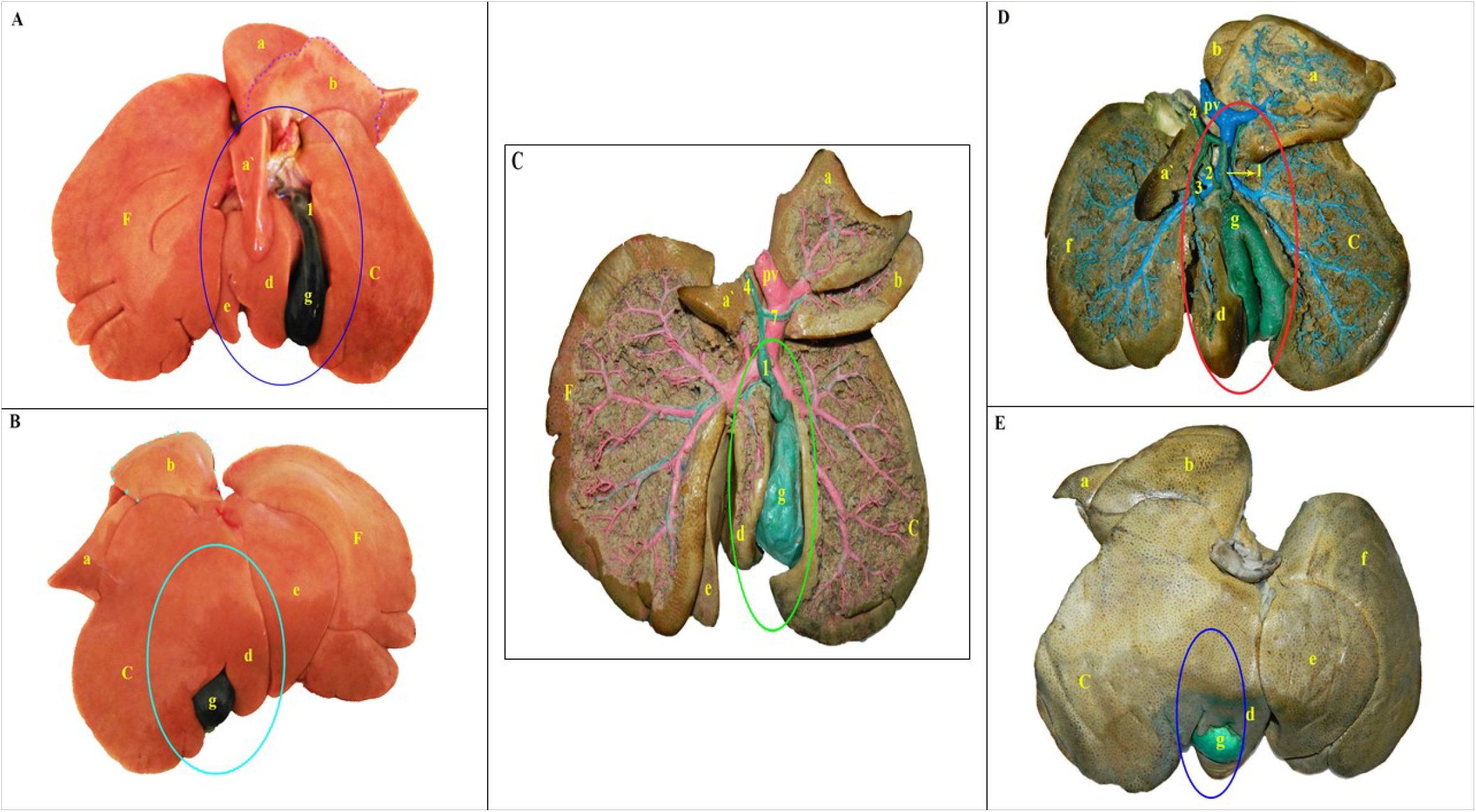
A photograph showing different shapes of gall bladder in cat. (A,B): Broad rounded fundus with wide body and narrow neck on visceral surface (A), on parietal surface (B), (C): Truncated fundus gall bladder, (D,E): Bilobed gall bladder on the visceral surface (D), on the parietal surface of the liver (E).

**Fig. (9):**
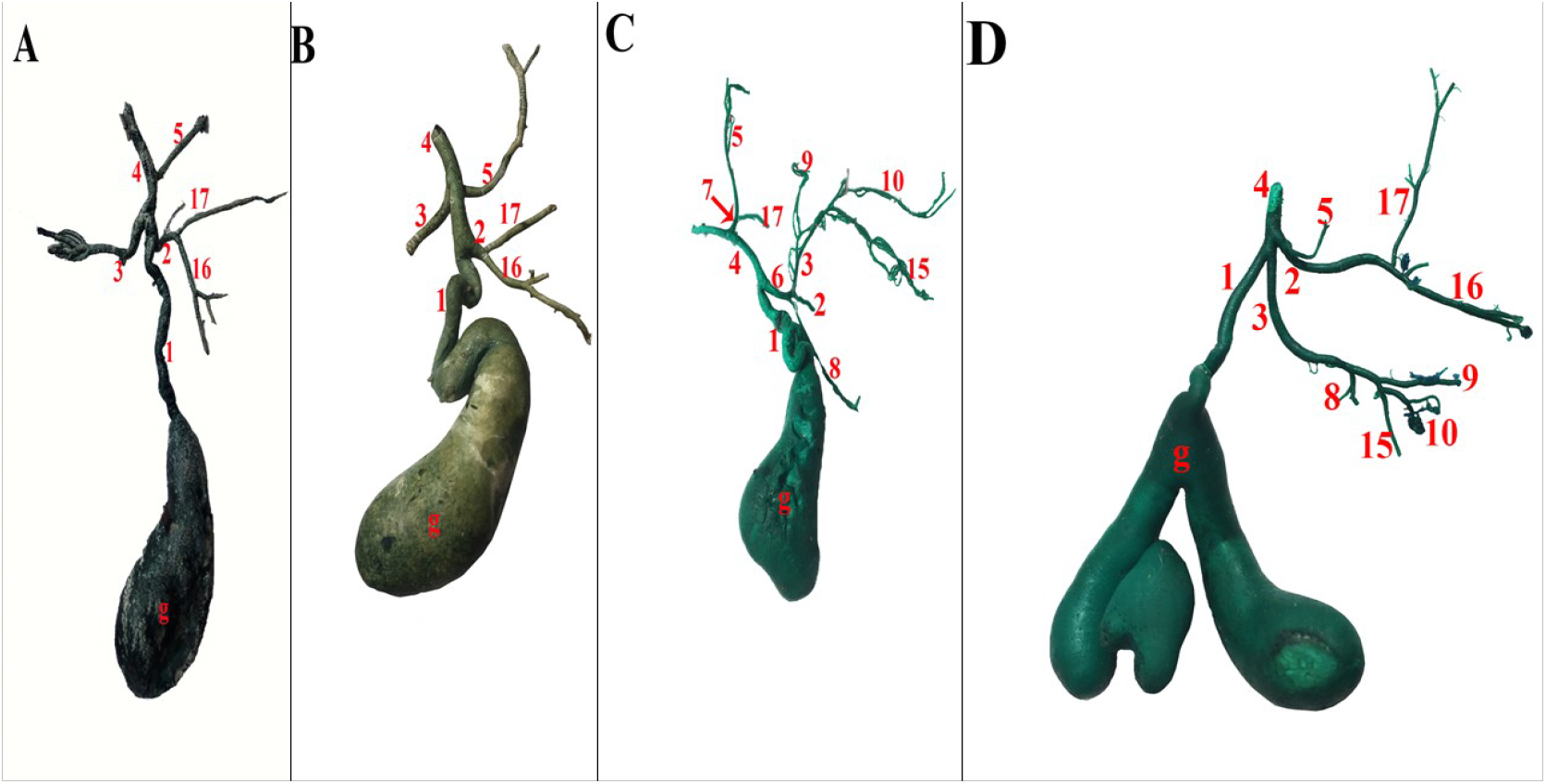
Latex neoprene cast of the cat biliary system isolated; (A,B,D): Bile duct formed by Triple conversion of the left and the right hepatic ducts with the cystic duct, (C): Bile duct formed by the union of the cystic duct with the common hepatic duct.

**Fig. (9):**
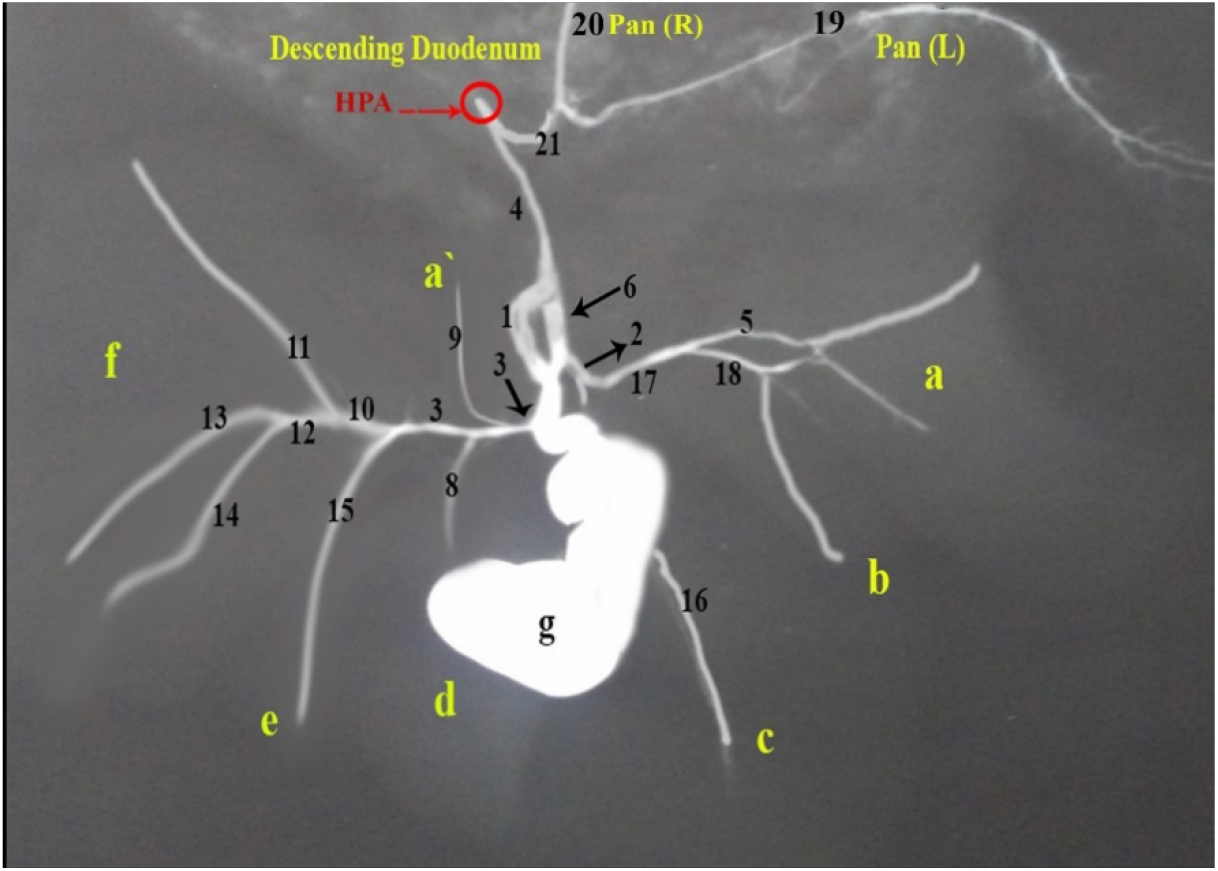
Latex neoprene cast of the cat biliary system isolated; (A,B,D): Bile duct formed by Triple conversion of the left and the right hepatic ducts with the cystic duct, (C): Bile duct formed by the union of the cystic duct with the common hepatic duct.

**Fig. (11):**
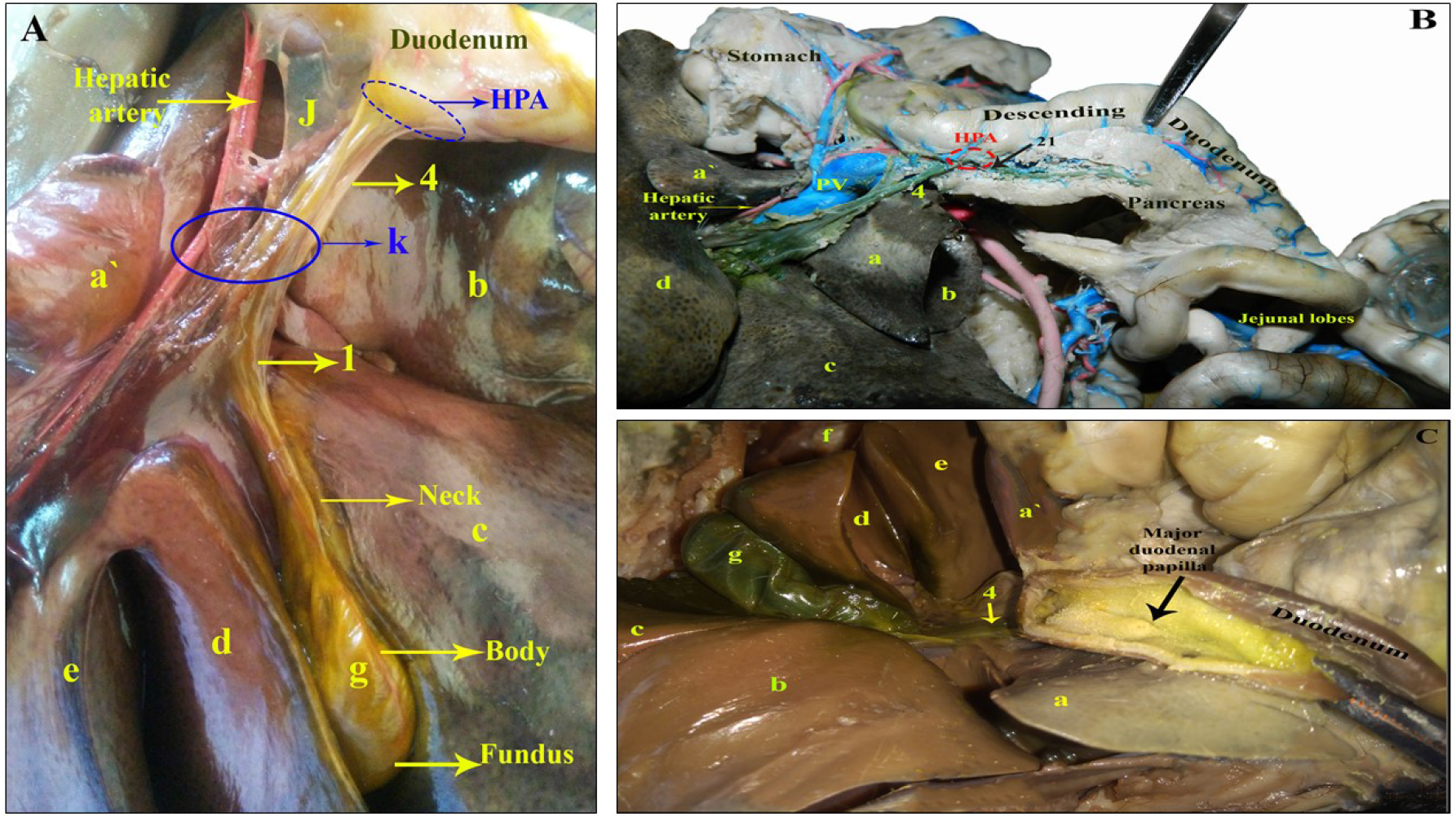
A photograph showing the course of the bile duct in the hepatoduodenal ligament (A), its union with the main pancreatic duct (B) and its opening on the major duodenal papilla (C) surrounded by the Hepatopancreatic ampulla in cat.

During the present investigation on cat liver, four different variations in its shape were observed;

**A**. The gall bladder was relatively small, partly appeared on the liver visceral surface, enveloped completely by the quadrate lobe and presented only a small impression on the right medial lobe. its broad fundus (Fig.7A/g) was divided by a sagittal groove (Duplex fundus) and each portion showed several small notches externally, appeared on the liver parietal surface (Fig.7C/g) as the gall bladder fossa in such case was too deep. Internally, there was no septa dividing the fundus or corpus (Fig.7B).
**B**. The gall bladder was fairly large and long to reach, nearly, the level of the liver ventral borders where they appeared on the parietal aspect (Fig. 8): **B.1**. The gall bladder fundus was broad and rounded (Figs. 8A,B, 9A/g), connected with wide body, gradually decreased to end by a narrow neck ventral to the hepatic porta. **B.2**. The gall bladder fundus appeared as it was truncated (Figs. 8C,9c/g) with slightly longer right end. The narrow-folded neck connected with a spiral cystic duct (Figs. 8C,9C/1). In such condition the gall bladder could be named (Truncated fundus). **B.3**. Bilobed gall bladder with duplex fundus, duplex corpus connected with single small neck (Figs. 8D,9D/g), tapered gradually and connected with a tortuous cystic duct. In such condition, the gall bladder neck and the proximal parts of the right and left lobes were bordered by the wide gall bladder fossa on the visceral surface of the liver, while the ventral portion of the right medial hepatic lobe embraced the distal half of right gall bladder lobe which carried two small buds (Fig. 9D/g) embedded in hepatic parenchyma of liver right lobe (Fig. 8D/g). The ventral portion of the gall bladder left lobe was partly covered by the quadrate lobe and its fundus appeared on the liver parietal surface (Fig. 8E/g).

The present investigation classified the formation of the feline bile duct and consequently the terminations of the hepatic ducts into two groups; **Group A;** The most common: The bile duct was formed by the triple convergence of the cystic duct with the left and right hepatic ducts (Figs.9A,B,D):

-The cystic duct (Figs. 9A,B,D/1) was relatively long and tortuous. The left hepatic duct (Figs. 9A,B,D/3) followed the left portal vein branch and formed by the lobar ducts from both left lobes and quadrate lobe and just before its termination, it received – in most cases – the papillary process branch (Fig. 9D/3,8,9,10,15). The right hepatic duct (Figs. 9A,B,D/2) was formed – commonly – by the junction of the right medial lobar duct (from the right medial lobe) and the right lateral lobar duct (drained the right lateral lobe as well as the caudate process by the Ramus processus caudatus) (Fig. 9D/2,5,16,17). It must be noted that the right hepatic duct in three specimens was only formed by the union of the right lateral and right medial lobar ducts which drained their respective lobes (Figs. 9A,B/2,16,17). In such conditions, the branch of the caudate process opened directly into the bile duct (Figs. 9A,B/4,5).

In **Group B:** It was revealed in two specimens in which the bile duct was formed by the union of the cystic duct with a short common hepatic duct (Figs. 9C,10):

-The cystic duct (Figs. 9C,10/1) was long and spiral. The common hepatic duct (Figs. 9C, 10/ 6) was short, formed by the union of the left hepatic duct collected both left lobes and papillary process (Fig. 9C/3,9,10,15) as well as receiving the duct of the quadrate lobe in one specimen (Fig. 10/8), while the right hepatic duct drained the right medial lobe branch and the quadrate lobar branch (Fig. 9C/2,8). In such specimens, the right lateral lobar branch and the branch of the caudate process united to form what could be called duct hepatica dextra accessoria (Fig. 9C/5,7,17) which opened directly into the bile duct. The latter detached a very thin ductule to the dorsolateral portion of right medial lobe.

#### Ductus choledochus

the cat bile duct (Figs. 10,11/4) was relatively long, proceeded through the hepatoduodenal ligament (Fig. 11A/J), where it merged with the pancreatic duct (Figs.10,11B/21) and pierced the duodenal wall to open on the major duodenal papilla surrounded by the hepatopancreatic ampulla (Figs. 10,11) 2-2.5 cm distal to pylorus.

### B. Ultrasonographical study

To facilitate the ultrasonographic examination of the abdominal organs, in the current work, we schemed the ventral abdominal surface of rabbit (Fig. 12I) and cat (Fig. 12II) according to the topographic anatomy. The abdomen was divided into three main regions (epi-, meso- and hypo-gastric regions) which were subdivided into three sub-regions forming a sum of nine abdominal sub-regions; the epigastric (Fig, 12/A) divided into left, right hypochondriac and middle xyphoid regions, the meso-gastric (Fig. 12/B) divided into left, right paralumbar (flank) and middle lumbar (umbilical) regions, as well as the hypogastric divided into left, right inguinal and middle pubic sub-regions. Our investigation of the gall bladder was mainly confined to the left medial hypochondriac, middle xyphoid and right medial hypochondriac epigastric region.

**Fig. (12):**
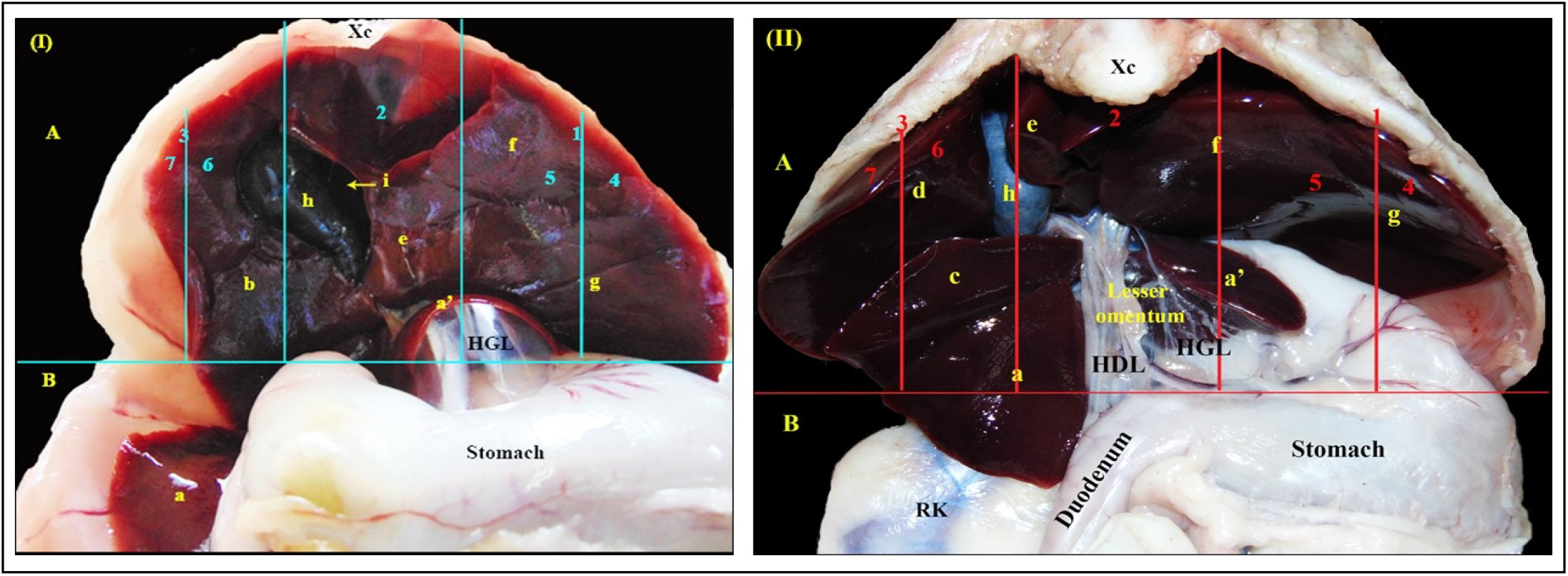
Photographs showing the topographic description of the abdominal epigastric and meso-gastric regions and the position of the liver in-situ in rabbit (I) and cat (II): A.Epigastric abdominal region, B.Meso-gastric abdominal region, 1.Left hypo-chondral epigastric region, 2.Middle (Xyphoid) epigastric region, 3.Right hypo-chondral region, 4.Left lateral hypo-chondral epigastric region, 5.Left medial hypo-chondral epigastric region, 6.Right medial hypo-chondral epigastric region, 7.Right lateral hypo-chondral epigastric region, a: Caudate process of caudate hepatic lobe, a’: Papillary process of caudate hepatic lobe, b: Right hepatic lobe, c: Right lateral hepatic lobe, d: Right medial hepatic lobe, e: quadrate hepatic lobe, f: Left medial hepatic lobe, g: Left lateral hepatic lobe, h: Gall bladder, HGL: Hepato-gastric ligament, i: fossa for gall bladder

The animals were positioned in dorsal recumbency and a 5-7.5 MHz and a 7.5 MHz linear multifrequency transducer was used for rabbit and cat, respectively. The sonographic approach was transabdominal percutaneous epigastric.

## DISCUSSION

### A. Anatomical study

The present study revealed that the rabbit gall bladder was relatively small, elongated in shape with a rounded fundus and narrow neck. It was positioned high in its fossa on the liver visceral surface between the right and the quadrate lobes and was fixed to the hepatic parenchyma by several small hepato-cystic ducts, thus it did not reach the liver ventral margin even when it was completely filled. In all investigated specimens, it did not appear on the liver parietal surface. A result which was in accordance with the description of **[7], [3] and [4]** in the same animal.

The results of our studies on the cat gall bladder revealed that in most specimens, it was relatively large, oval or slightly curved pear-shaped, situated more superficially in its fossa between the quadrate and right medial lobe and neither reach the liver ventral margin nor the parietal surface. In this respect, the canine gall bladder was embedded within the liver and appeared on the visceral and parietal surfaces **[32]; [10] and [18].**

On the other hand, our investigations on the same examined cat livers, revealed four different shapes and sizes of the gall bladders. In one case, the gall bladder was relatively small pear-shaped with duplex fundus, without any internal septa. In second case, the liver gall bladder was bilobed; had duplex fundusm duplex corpus with single small neck. The previous results were reported as congenital abnormalities by **[21]; [22] and [23].**

Moreover, a third investigated cat liver showed a distended long gall bladder with rounded fundus reaching the ventral margin of the liver and appearing from its parietal surface. A result which simulated that revealed by **[24]** in anorexic cat. The fourth exceptional specimen in cat was long gall bladder, reaching the ventral border with slightly larger right end and could be called truncated fundus.

It must be noted that, the liver investigated specimen of bilobed gall bladder revealed that the right lobe presented two small buds connected together. A condition which was not recorded in relevant references. Our results on cat liver was documented by **[20]** in that the variety of the cat gall bladder shape vacuolization of the solid gall bladder and bile duct diverticulum during development.

Concerning the extrahepatic biliary tract, the most common pattern of rabbit bile duct formation investigated in the present study, by the junction of the left hepatic duct with the cystic duct. The left duct drained both left lobes and quadrate lobe as well as the papillary process of the caudate lobe, while the right hepatic lobe was drained by its own right hepatic duct which opened into the cystic duct about its middle in its route to enter in the formation of the bile duct. The bile of the caudate process of the caudate lobe was collected by two branches that merged and opened directly into the bile duct.

The absence of the common hepatic duct in the rabbit most examined specimens was in accordance with the description of **[3] and [4]** in the same animal as well as **[5] and [6]** in guinea pig. On the other hand, the statement of **[7]** that the formation of the bile duct by joining the common hepatic duct with the cystic duct was observed only in two out of six rabbit liver specimens where the cystic duct was very thin and joined the fairly large right hepatic duct. The latter merged with the left duct to form a short common hepatic duct that continued in the hepatoduodenal ligament as the bile duct.

The present investigation on the most examined cat liver revealed the formation of the bile duct by the triple convergence of the left, right hepatic and cystic ducts. A result which was confirmed by the document of **[12]** in the same animal. Moreover, the present work recorded, in two specimens, that the right hepatic duct joined with the left one to form a short common hepatic duct that merged the cystic duct to form the bile duct. In this respect, the absence of the common hepatic duct and the junction of the lobar hepatic ducts individually with the cystic duct to form the bile duct as described in canine by **[8], [33], [34], [32] and [3]** as well as **[11]** was in contrary to our investigations.

In the rabbit investigated specimens, the caudate process of the caudate lobe was drained by its own branch that entered directly into the bile duct. A result which was in agreement with **[7]** in the same animal and also in ruminants **[13]; [14] and [35]**.

The biliary ducts drained the caudate and the right lateral lobes entered the bile duct directly was observed by **[9]** in prairie dog and in domestic animals **[18]**. Similar observation was also recorded in the present study on some cat livers. However, in most examined specimens the two ducts merged to form what could be named right accessory hepatic duct. However, the present study revealed, in exceptional cases, that caudate process branch entered separately the bile duct while the right lateral lobar branch entered in the formation of the right hepatic duct by joining the right medial lobar branch.

The entrance of the bile and pancreatic ducts into the duodenum either individually as in the investigated rabbit specimens or ended together as observed in the cat specimens were in accordance with **[7] and [3]** in the rabbit as well as in the cat **[33]; [3]; 1 and [36]**.

Regarding the presence of the hepatopancreatic ampulla surrounding the common orifice of the bile and pancreatic ducts observed in the investigated cat specimens, was in agreement with **[33] and [37]** in the same animals as well as in the guinea pig **[19]**.

### B. Ultrasonographical study

The ultrasonographic study on the rabbit’s liver was performed using a linear multifrequency transducer with a working frequency range from 5 to 7.5 MHz, following **[38].** This contrasted the opinion offered by **[39]** who used a micro convex probe within the same range of frequency. In cats, a linear multifrequency transducer with working frequency 7.5 MHz was used to scan cat’s liver in accordance with **[40]** and nearly agreed with **[41] and [42]** who also declared using the same range of frequency but with a curved array transducer for abdominal ultrasonography in cats. Meanwhile, **[43]** offered using a 5 MHz transducer.

Our examination on the liver was performed while the animals in dorsal recumbency as asserted by **[43]; [44] and [45]** in cats and **[39] and [38]** in rabbits. However, other authors recommended to use a combination of positions; right lateral, left lateral and dorsal recumbency **[46] and [47]** and **[46]** who added that the gall bladder was better detected with the animal in lateral recumbency.

In the current study, the ultrasonographic approach was transabdominal percutaneous epigastric in both rabbits and cats, this approach was in consent to [39] **and [38]**. However, the liver was observed from a ventral abdominal approach **[45]** or by placing the transducer just caudal to the xiphisternum **[43] and [45]** or beneath the costal arch **[48].** All whoever agreed that the liver could be accessible unlike what was proposed that the ultrasonographic examination of the liver in the cat is difficult due its location under the rib cage **[40]** or a portion of the liver being obscured by the overlying stomach **[48].**

The liver could be detected successfully on sonar by identifying the neighboring anatomic landmarks; different soft and hard tissues were identified including the hyperechoic diaphragm cranially and the stomach caudally as agreed with **[44], [48] and [47].** In addition to the costal arch laterally, the xyphoid cartilage ventrally as well as the right kidney caudally, this statement was in line with the findings of **[39].**

In rabbit, the right medial hypo-chondral sagittal scan, identifying the hypoechoic right hepatic lobe could be confirmed by the passage of the caudal vena cava dorsally from the thoracic cavity till perforating the diaphragm at the level of caval foramen. The caudal vena cava appeared tubular anechoic with no visible wall as agreed by **[49]** and the small, slightly elongated gall bladder with rounded fundus could also be seen, which appeared with anechoic lumen and no visible wall. Furthermore, the middle (xyphoid region) sagittal scan showed the xyphoid cartilage entirely masking the underlying middle hepatic lobes accompanied by distal acoustic shadowing, but during the examination, the pressure was applied under the xyphoid and the left medial, quadrate and right hepatic lobes were identified respectively from left to right direction and the anechoic gall bladder appeared with no visible wall which was found between the quadrate hepatic lobe on the left and the right hepatic lobe on the right.

In consent to the recommendation of **[38]**, the rabbit’s liver was evaluated in both longitudinal and transverse scanning planes. In this regard, the right medial hypo-chondral transverse scan showed hypoechoic right and quadrate lobes between which lies the anechoic gall bladder with no visible wall, normally accompanied by mirror and distal acoustic enhancement artifacts. Moreover, in the middle (xyphoid region) transverse scan, the characteristically broad oval-shaped xyphoid cartilage entirely masked the middle lobes of the liver and was normally accompanied by distal acoustic shadowing, but with applied pressure under the xyphoid cartilage, hypoechoic left medial, quadrate and right lobes appeared, respectively from left to right direction. Also, the elongated oval gall bladder appeared anechoic with hypoechoic wall lying between the quadrate (left) and right lobe (right) from which emerging the cystic duct as detected by **[39].**

In the present investigation in cat, in the right medial hypo-chondral transverse scan, the identification of the anechoic gall bladder with distinct echogenic wall contributed to the determination of the two bordering lobes; the quadrate and the right medial lobes. The observation of the gall bladder was in accordance to that of **[43]; [46]; [45] and [42]** but was not recorded in the statement of **[48] and [50]** who announced that the gall bladder wall was poorly visualized or not identified at all. Incoherence to the anatomic description of the feline gall bladder, it varied considerably in shape and size as approved by **[48].**

In the left hypo-chondral sagittal scan -in case of an extended stomach – no trace of the gall bladder was seen, only the left lateral and the left medial lobes appeared with the c/s of V. hepatica lobi sinistri lateralis branches having anechoic lumen without a distinct echogenic wall. while in case of an empty stomach, only the hypoechoic quadrate lobe appeared as well as the pear-shaped anechoic gall bladder surrounded by echogenic wall.

Having the opinion that the intrahepatic bile ducts could not be evident on a general ultrasonographic examination in normal cats, our examination was the same in both examined animals; rabbit and cat, as that demonstrated by **[43]; [40]; [48]; [45]; [47]**. However, **[43]** offered that the extrahepatic bile duct could be visualized as offered in our observation of the cystic duct in rabbit in the middle transverse scan.

### CONCLUSION

The rabbit gall bladder was relatively small in all investigated liver specimens. It was positioned high and fixed by several small hepato-cystic ducts to its fossa. The rabbit bile duct was formed commonly by the junction of the left hepatic duct and the cystic duct. The former one drained both left lobes and quadrate lobe as well as the papillary process of the caudate lobe. the cystic duct was, commonly fairly large, received the right hepatic duct that collected the right lobe in its route to enter the duodenum, the bile duct receives the branch of the caudate process of the caudate lobe. It must be noted that, in some of the investigated specimens, the rabbit gall bladder was connected by a thin and small cystic duct, with the right hepatic duct which joined the left one to form a short common hepatic duct.

The relatively large, oval or slightly curved pear-shaped cat gall bladder were commonly investigated. It was situated more superficially on the middle of the liver visceral surface and neither reached the ventral margin nor appeared on its parietal surface. In addition, the present study revealed other four anatomic variations dealing with the shape and size of the feline, native breed, gall bladder from fundic duplication, fundic duplication with body duplication (Bilobed), truncated fundus and distended rounded fundus. In all these different forms, the gall bladder appeared partially on the hepatic diaphragmatic surface. There was no relation between the shape of the gall bladder and the mode of the hepatic ducts’ terminations in felines. Commonly, the bile duct was formed by the triple convergence of the left and the right hepatic ducts with the cystic duct. However, in some exceptional cases a short common hepatic duct was formed, by the union of the previous two hepatic ducts, and merged with the cystic duct forming the bile duct. In the present study, the caudate process branch and the right lateral branch, each drained its respective hepatic portion. Commonly, these two branches merged to form a single duct that could be named right accessory hepatic duct which entered the “original” right hepatic duct of the right medial lobe.

The sonographic investigation of the gall bladder was mainly confined to the left medial hypochondriac, middle xyphoid and right medial hypochondriac epigastric region. The normal gall bladder in rabbit appeared small, elongated with anechoic lumen bordered by right lobe laterally and quadrate lobe medially and has no visible wall, but in cat varied in conformation, bordered by the right medial lobe laterally and the quadrate lobe medially surrounded by echogenic wall.

## ACKNOWLEDGMENT

We would like to express our great appreciation to professor doctor **Salah Mohamed Hagrass**; professor of Anatomy and Embryology, faculty of veterinary medicine, Cairo University, for his valuable and constructive work during the development of this research work. His willingness to give his time so generously has been very much appreciated.

## Legends for figs. (1 – 4) in Rabbit

**a. Processus caudatus; Lobus caudatus, a’.Processus papillaris; Lobus caudatus, b. Lobus hepaticus dexter, c. Lobus quadratus, d. Lobus hepaticus sinister medialis, e. Lobus hepaticus sinister lateralis, f. Vesica fellea, g. Lesser omentum, h. hepatoduodenal ligament, 1. Ductus choledochus, 2. Ductus hepaticus communis, 3. Ductus hepaticus sinister, 4. Ductus hepaticus dexter, 5. Ductus cysticus, 6. R. lobi dexter, 7. Ramus lobi sinistri laterals, 8. R. dorsalis lobi sinistri lateralis, 9. R. intermedius lobi sinistri lateralis, 10. R. ventralis lobi sinistri lateralis, 11. R. lobi sinistri medialis, 12. R. medialis lobi sinistri medialis, 13. R. lateralis lobi sinistri medialis, 14. R. lobi quadratus, 15. R. omentalis (R. processus papillaris), 16. R. Processus caudatus, 17. R. Visceralis processus caudatus, 18. R. Parietalis processus caudatus, 19.Rr. quadrati, 20. Rr. Ventralis lobi dexter, 21. Rr. Dorsalis lobi quadrati.**

## Legends for figs. (5 – 11) in Cat

**a. Processus caudatus; Lobus caudatus, a’.Processus papillaris; Lobus caudatus, b. Lobus hepaticus dexter lateralis, c. Lobus hepaticus dexter medialis, d. Lobus quadratus, e. Lobus hepaticus sinister medialis, f. Lobus hepaticus sinister lateralis, g. Vesica fellea, h. Lesser omentum, i. Hepatogastric ligament, J. Hepatoduodenal ligament, k. Porta hepatis, HPA: Hepatopancreatic ampulla, Pv: Portal vein, Pan (R): Right pancreatic lobe, Pan (L): Left pancreatic lobe, 1. Ductus cysticus, 2. Ductus hepaticus dexter, 3. Ductus hepaticus sinister, 4. Ductus choledochus, 5. Ramus processus caudatus, 6. Ductus hepaticus communis, 7. Ductus hepaticus dextra accessoria, 8. Rr. lobi quadrati, 9. R. omentalis, 10. Ramus lobi sinistri laterals, 11. R. dorsalis lobi sinistri lateralis, 12. Truncus communis, 13. R. intermedius lobi sinistri lateralis, 14. R. ventralis lobi sinistri lateralis, 15. R. lobi sinistri medialis, 16. R. lobi dexter medialis, 17. R. lobi dexter lateralis, 18. Rr. lobi dexter lateralis, 19. Left pancreatic duct, 20. Right pancreatic duct, 21. Main pancreatic duct.**

**Fig. (13):**
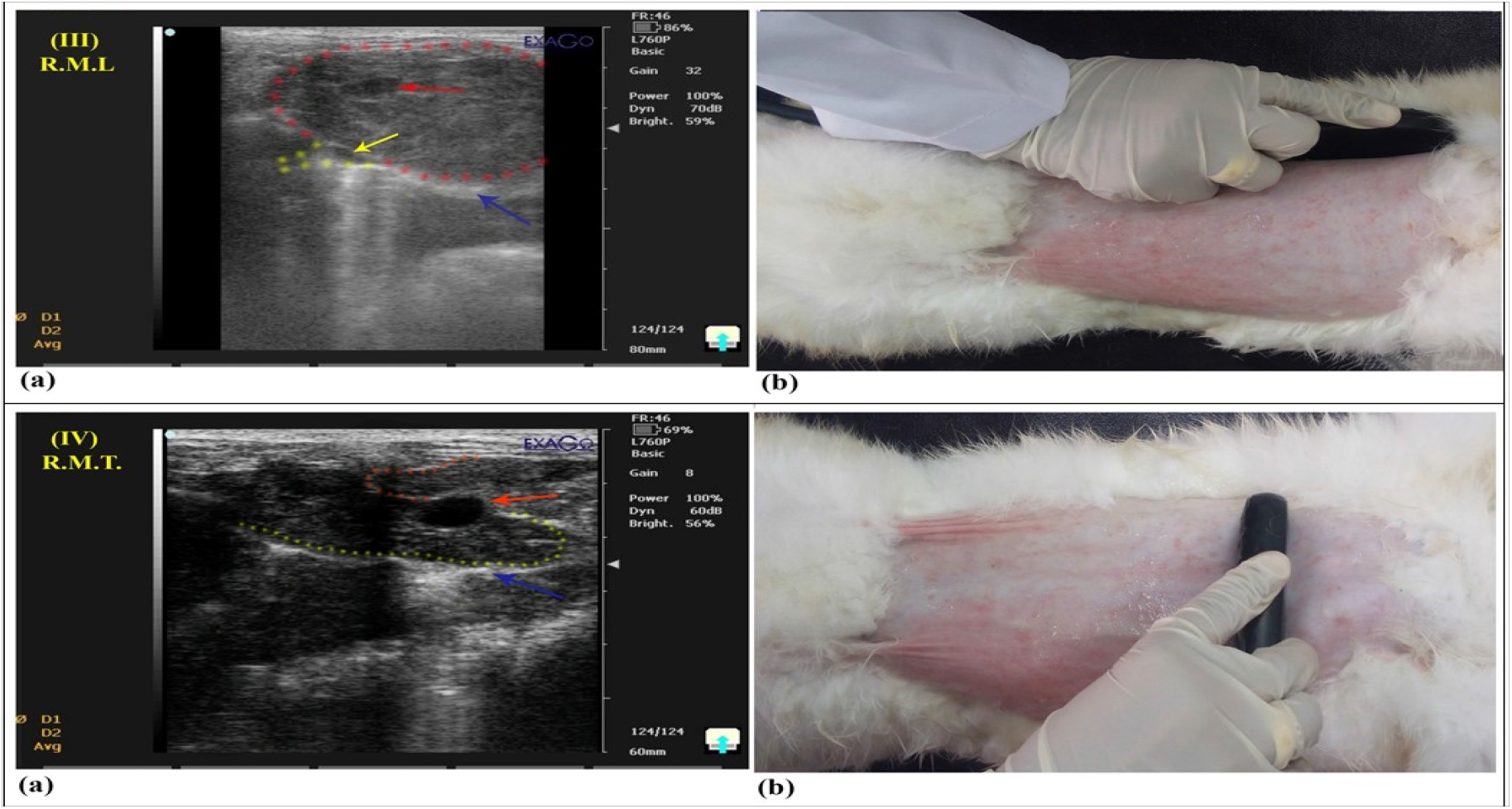
**A photograph showing the right medial hypo-chondral epigastric ultrasound scan in rabbit in sagittal plane (III) and transverse plane (IV), (III) Right Medial Longitudinal (RML) ultrasound scan showing;** a.Hypoechoic Right lobe outlined by red dots and a red arrow pointing at the small, slightly elongated gall bladder with rounded fundus, which appeared with anechoic lumen with no visible wall. Noted yellow dots showing the caudal vena cava appearing tubular anechoic lumen and no visible wall emerging from the thoracic cavity into the echogenic diaphragm (blue arrow) at the level of caval foramen (yellow arrow). b.Location and direction of linear transducer (Right medial hypo-chondral epigastric longitudinal). **(IV) Right Medial Transverse (RMT) ultrasound scan showing;** a.Hypoechoic Right and Quadrate lobes outlined by yellow and red dots, respectively-with a red arrow pointing at the anechoic gall bladder having no visible wall and a blue one pointing at the echogenic diaphragm. Noted the normally accompanying Mirror and distal acoustic enhancement artefacts. b.Location and direction of linear transducer (Right medial hypo-chondral epigastric transverse).

**Fig. (14):**
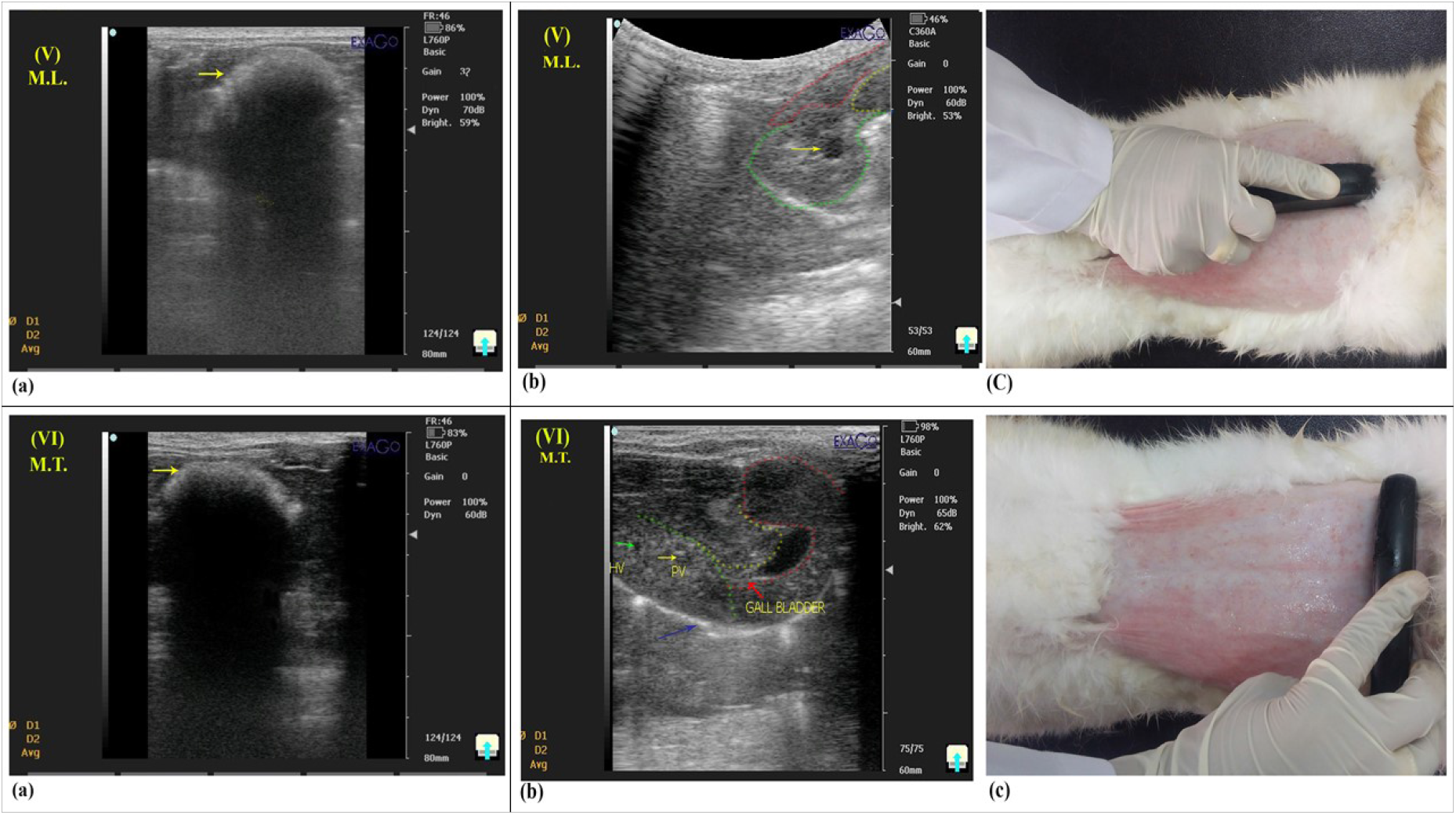
**A photograph showing the middle xyphoid epigastric ultrasound scan in rabbit in sagittal plane (V) and transverse plane (VI), (V) Middle Longitudinal (ML) ultrasound scan showing;** a.The xyphoid cartilage (yellow arrow) completely masking the middle lobes of the liver and normally accompanied by distal acoustic shadowing. b.With pressure under the xyphoid, hypoechoic left medial, quadrate and right lobes appeared and outlined by red, yellow and green dots, respectively. Noted the yellow arrow pointed at the gall bladder which appeared anechoic with no visible wall on the middle of the liver visceral surface between the quadrate lobe medially and right lobe laterally. c.Location and direction of the linear transducer (Middle xyphoid epigastric longitudinal). **(VI) Middle Transverse (MT) ultrasound scan showing;** a.The same as (V)a. b.With pressure under the Xyphoid, hypoechoic left medial, quadrate and right lobes appeared and outlined by green, yellow and red dots, respectively-with characteristic C/S appearance of anechoic vena hepatica lobi sinistri medialis without echogenic wall (green arrow) and anechoic ramus lobi sinistri medialis portal vein with echogenic wall (yellow arrow). Noted the small elongated anechoic gall bladder with a red arrow pointed at the cystic duct and a blue arrow pointed at the echogenic diaphragm. c.Location and direction of linear transducer (Middle xyphoid epigastric transverse).

**Fig. (15):**
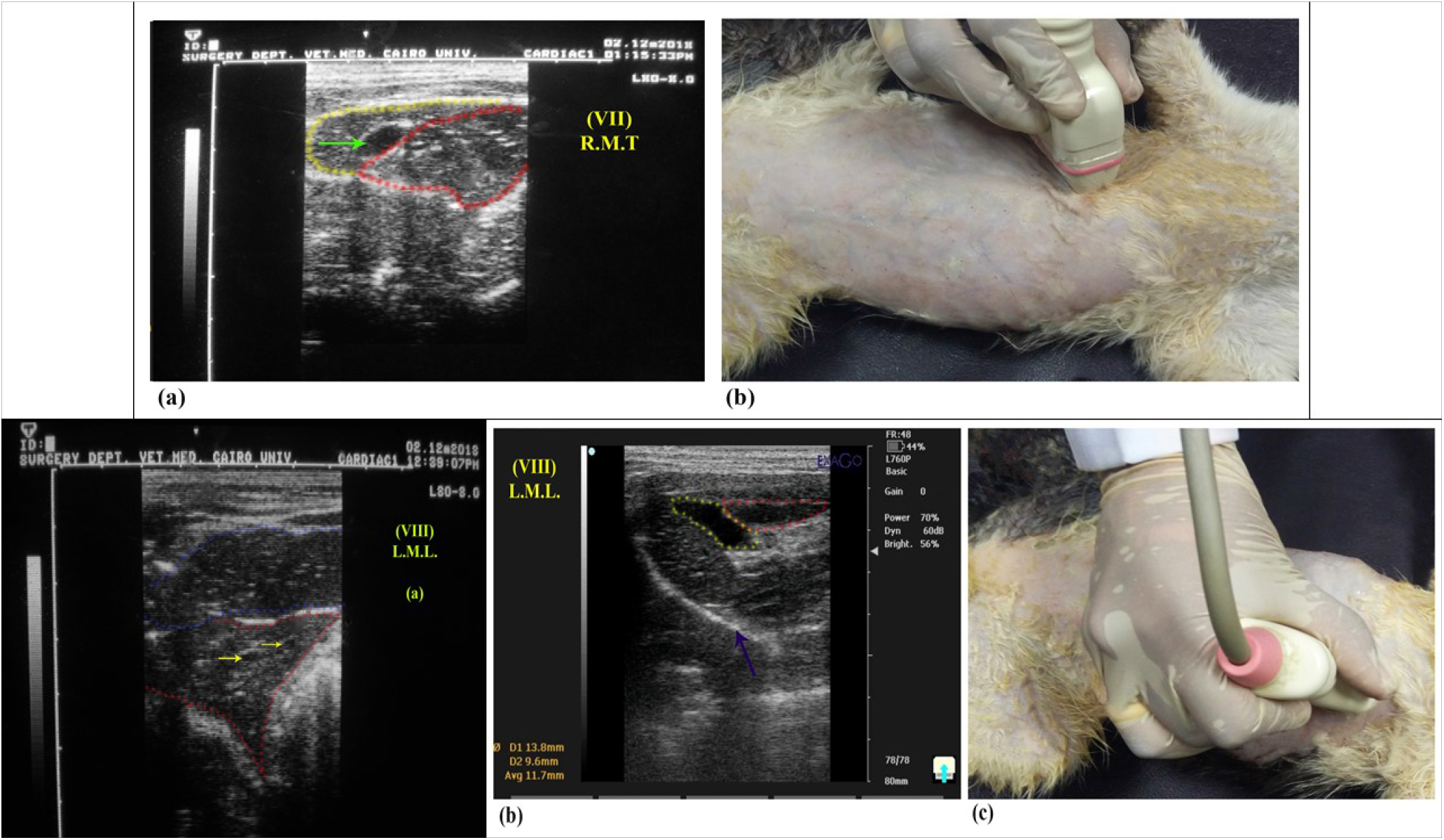
**A photograph showing the right and left medial hypo-chondral epigastric ultrasound scan in cat in transverse (VII) and sagittal planes (VIII), respectively (VII) Right Medial Transverse (RMT) ultrasound scan showing; a.**Hypoechoic Right medial (red dots) and quadrate (yellow dots) lobes with a green arrow pointing at the gall bladder appearing with anechoic lumen surrounded by echogenic wall, situated between them. **b.**Location and direction of linear transducer (Right medial hypo-chondral epigastric transverse). **(VIII) Left Medial Longitudinal ultrasound scan showing; a.**In case of extended stomach, no trace of gall bladder was observed, only the hypoechoic heterogenous left lateral and left medial lobes appeared outlined by red and blue dots, respectively. The yellow arrows are pointed at the C/S of the branches of v. hepatica lobi sinistri lateralis appearing anechoic lumen without echogenic wall. **b.**In case of empty stomach, the pear-shaped anechoic gall bladder (yellow dots) appeared surrounded by echogenic wall in close relation to only the hypoechoic quadrate lobe appeared outlined by red dots. Noted the blue arrow pointed at the echogenic diaphragm. **c.**Location and direction of linear transducer (Left medial hypo-chondral epigastric longitudinal).

